# High behavioural variability mediated by altered neuronal excitability in *auts2* mutant zebrafish

**DOI:** 10.1101/2020.11.04.367821

**Authors:** Urvashi Jha, Igor Kondrychyn, Vladimir Korzh, Vatsala Thirumalai

## Abstract

Autism spectrum disorders (ASDs) are characterized by abnormal behavioral traits arising from neural circuit dysfunction. While a number of genes have been implicated in ASDs, in most cases, a clear understanding of how mutations in these genes lead to circuit dysfunction and behavioral abnormality is absent. The *autism susceptibility candidate 2* (*AUTS2*) gene is one such gene, associated with ASDs, intellectual disability and a range of other neurodevelopmental conditions. Yet, the function of AUTS2 in neural development and circuit function is not at all known. Here, we undertook functional analysis of Auts2a, the main homolog of AUTS2 in zebrafish, in the context of the escape behavior. Escape behavior in wild type zebrafish is critical for survival and is therefore, reliable, rapid, and has well-defined kinematic properties. *Auts2a*^−/−^ zebrafish are viable, have normal gross morphology and can generate escape behavior with normal kinematics. However, the behavior is unreliable and delayed, with high trial-to-trial variability in the latency. We demonstrate that this is due to the reduced excitability of Mauthner neurons resulting in unreliable firing with stimuli that normally elicit the escape response. Combined with previous studies that show Auts2-regulation of the transcription of ion channel proteins, our results suggest that Auts2 sets the excitability of neurons by activating a set transcriptional program.

**Significance statement:** AUTS2 is one among recently identified autism susceptibility candidate genes, whose function in neuronal circuits is unclear. Using zebrafish as a model organism, we probe the function of Auts2a (homolog of mammalian AUTS2) at the cellular, network and behavioral levels. The escape behavior of Auts2a mutant zebrafish is highly variable with normal short latency escapes, long latency escapes and total failures across trials in the same fish. This occurs because neuronal excitability is inappropriately set in the Mauthner neurons of mutants leading to the large trial-to-trial variability in responses. The behavioral variability is fully explained by variability in firing action potentials in the Mauthner neuron, providing an integrative understanding of how behavioral variability arises from mutations at the genetic level.

## Introduction

Every neuron needs to carefully tune its excitability to be able to perform computation within the circuit in which it is embedded. When firing properties are improperly specified, neuronal function is compromised leading to abnormal behaviors. Transcriptional regulation plays important roles in specifying neuronal excitability properties and therefore, are also a chief class of genes implicated in neurological diseases such as Autism spectrum disorders (ASDs; (Bourgeron, 2015; De Rubeis et al., 2014; Sztainberg and Zoghbi, 2016)). The *autism susceptibility candidate 2* (*AUTS2*, also known as activator of transcription and developmental regulator) gene is a known regulator of transcription in the nervous system (Gao et al., 2014; Russo et al., 2018; Wang et al., 2018) and is associated with several neurodevelopmental disorders including ASD (Beunders et al., 2013; Kalscheuer et al., 2007; Sultana et al., 2002). AUTS2 is expressed in neurons of the central nervous system and is present both in the nucleus and in the cytoplasm (Bedogni et al., 2010; Gao et al., 2014; Hori et al., 2014; Hori and Hoshino, 2017). In the nucleus, AUTS2 binds to members of the Polycomb Repressor Complex 1 (PRC1), but activates transcription of several genes important for neural development and function (Gao et al., 2014; Oksenberg et al., 2014; Wang et al., 2018). In the cytoplasm, AUTS2 regulates actin cytoskeleton to control neuronal migration and neurite outgrowth (Hori et al., 2014; Hori and Hoshino, 2017). *Auts2* knockout-mice exhibited several deficits such as reduced righting reflexes and ultrasonic vocalizations (Gao et al., 2014). Nevertheless, how AUTS2 controls nervous system development, function and behavioral output are not understood at all.

We identified and characterized four paralogs of *auts2* in zebrafish: *auts2a*, *auts2b*, *fibrosin-like 1*(*fbrsl1*) and *fibrosin* (*fbrs*) (Kondrychyn et al., 2017). Both *auts2a* and *auts2b* genes are expressed in the developing and juvenile zebrafish brain. Analysis of gene structure and protein sequence revealed that among the two genes, *auts2a* is the closest orthologue to mammalian *Auts2* (61.58% identity in protein sequence), with even higher homology in the C-terminus, hinting at conserved binding partners and function. In larval zebrafish, *auts2a* is widely expressed in the brain with distinctly high expression in rhombomere 4 (Kondrychyn et al., 2017), which houses neurons of the escape network (Metcalfe et al., 1986).

Teleost escape behavior consists of a sharp C-shaped tail bend away from the inducing stimulus with only a few milliseconds latency (Kimmel et al., 1974; Eaton et al., 1977). The behavior is triggered by action potential firing in one of two bilaterally located giant Mauthner neurons (M-cells) (Eaton and Farley, 1975; Eaton et al., 1977; Korn and Faber, 2005; Kohashi and Oda, 2008; Sillar, 2009). Two pairs of homologous neurons, MiD2cm and MiD3cm, also take part in escape behaviors but fire at much longer latencies (Eaton et al., 1984; Kohashi and Oda, 2008).

Action potential firing in M-cells is required for the fast C-start response (Zottoli, 1977; Eaton et al., 1981). In response to supra-threshold depolarization via strong synaptic inputs or by direct current injection, M-cells in larval and adult zebrafish, as well as adult gold fish, generate a single action potential with a very short latency (Eaton et al., 2001; Nakayama and Oda, 2004; Watanabe et al., 2013). This action potential is conducted quickly via its giant axons to spinal circuitry, including direct synapses onto contralateral motor neurons, resulting in rapid muscle contraction and a sharp bending of the body (Fetcho, 1991). M-cell excitability is thus critical for quick escape from threatening stimuli. Though not all of the conductances driving M-cell response have been delineated, it is clear that during development, M-cell intrinsic properties are progressively tuned to result in its mature firing behavior (Brewster and Ali, 2013; Watanabe et al., 2017, 2013). Since *auts2a* is expressed in rhombomere 4 at stages when M-cell properties are being defined, we sought to determine how Auts2a impacts excitability of M-cells and therefore the escape behavior itself.

Using customized transcription activator-like effector nucleases (TALENs), we generated mutations in the *auts2a* locus and isolated an allele which had a premature stop codon in the coding sequence. Using a combination of high speed videography and *in vivo* calcium imaging, we show that escape behaviors become highly unreliable and slow in *auts2a* mutants and that this unreliability can be explained by the reduced excitability of M-cells. These results indicate a role for Auts2a in regulating neuronal excitability, an action by which Auts2a impacts behaviors significantly.

## Results

### Generation of the *auts2a* knockout zebrafish line

A pair of TALENs were designed to target the donor splice site at exon 8 of the *auts2a* gene (Figure 1A). The TALEN-targeted sequences surround a restriction enzyme site for easy screening through introduction of a restriction fragment length polymorphism. We identified 5 different mutant alleles, 3 of which led to a frameshift after S498 and premature stop codons after several missense amino acids (Figure 1B) and one of them, *auts2a*^*ncb104*^, was selected to establish an *auts2a* KO zebrafish line. This line harbours a 11-nt deletion, which disrupts the donor splice site affecting correct splicing between exon 8 and exon 9. RT-PCR analysis of RNA isolated from homozygotes revealed, that *auts2a*^*ncb104*^ pre-mRNA uses two alternative cryptic donor splice sites found in intron in order to splice exon 8 to exon 9 (Figure 1A, B). As a result, intronic sequence is partially retained in *auts2a*^*ncb104*^ mRNA leading to a premature stop codon at amino acid 504 after 6 missense amino acids (Figure 1B).

**Figure 1.**
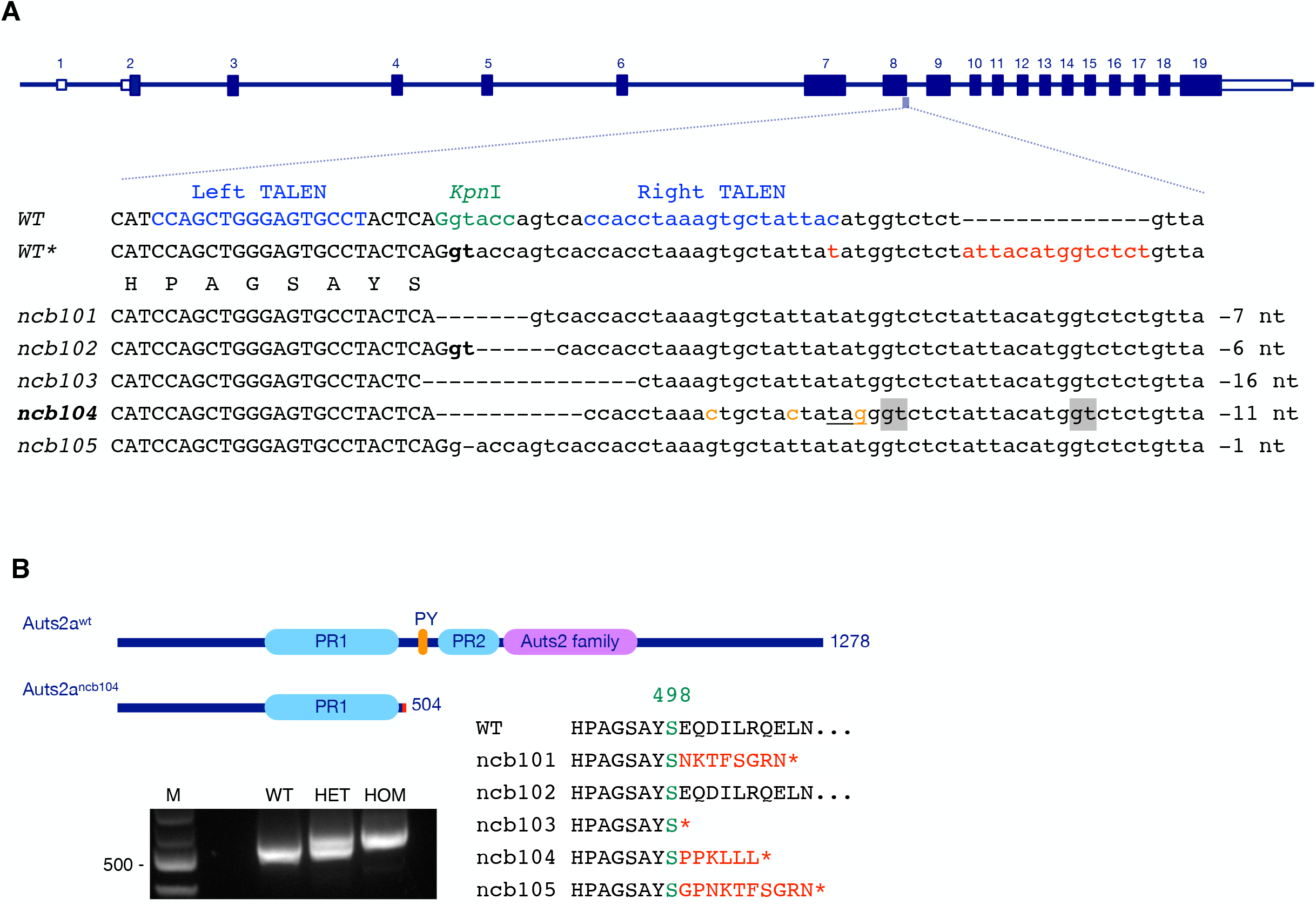
TALEN-induced mutation in *auts2a* gene. **(A)** Zebrafish *auts2a* gene locus, TALEN target sites and isolated alleles. The TALENs target a pair of binding sites (in blue) flanking a spacer with a restriction enzyme site (in green). Exonic and intronic sequences are shown in upper and lower cases, respectively. In contrast to genomic sequence annotated in the Ensembl (WT), our “in-house” zebrafish AB strain (WT*) has a polymorphism in intronic sequence adjacent to exon 8 (in red). Alleles *ncb10*1, *ncb103*, *ncb104* and *ncb105* have the nucleotide deletions that disrupt the donor splice site (in bold) leading to a frameshift after S498 and premature stop codons. Deletion in allele *ncb102* does not affect a correct splicing and Auts2 protein sequence. Three single point mutations were introduced in intron of *auts2a*^*ncb104*^ allele (in orange) leading to a stop codon creation (underlined). The alternative donor splice sites used to splice a mutant *auts2a*^*ncb104*^ pre-mRNA are highlighted in grey. **(B, top)** Auts2a and Auts2a^ncb104^ proteins. The *ncb104* mutation causes the loss of the C-terminal portion of Auts2a, comprising PY motif, proline-rich region PR2 and the Auts2 family domain. **(B, bottom left)** RT-PCR analysis of *auts2a* mRNA, isolated from wild type (WT), *ncb104* heterozygote (HET) and homozygote (MUT) embryos. M, 100-bp DNA ladder (NEB). **(B, bottom right)** Partial protein sequences of mutant alleles.

The zebrafish Auts2a protein has several domains, previously predicted in human AUTS2 (Sultana et al., 2002): two proline-rich (PR) regions, PR1 at amino acids 273-492 and PR2 at amino acids 558-656, PY (PPPY) motif at amino acids 524-528, and the Auts2 family domain at amino acids 660-882 (Figure 1B). In Auts2a^ncb104^, only the PR1 region is retained (Figure 1B). Recently, we described the transcriptional complexity found in the zebrafish *auts2a* gene locus that is mediated by alternative splicing and alternative promoter usage (Kondrychyn et al., 2017). Exon 8 is a common exon in all isoforms (Supplemental Figure S1A) and in *auts2a*^*ncb104*^ mutant all isoforms will be similar affected: a loss of the C-terminal portion of Auts2a, comprising PY motif, PR2 region and the Auts2 family domain (Supplemental Figure S1B).

### *auts2a* mutants display high variability in escape responses

*Auts2a* KO zebrafish showed normal development and gross morphology (Supplemental Figure S2A). Moreover, *auts2a*^*ncb104*^ homozygote fish survive to become fertile adults. As *auts2a* is a neurodevelopmental gene (Oksenberg and Ahituv, 2013) and is expressed at very high levels in rhombomere 4 (Kondrychyn et al., 2017), which houses the escape network, we first investigated whether M-cells, the command-like neurons driving escape behavior, are present in *auts2a* mutants. Wild type larvae possess a single pair of M-cells that send commissural axons down the spinal cord (Fetcho, 1991). In *auts2a* mutants both M-cells are present and send commissural axons (Supplemental Figure S2B). However, M-cells in *auts2a* mutants were smaller: both soma volume and dendritic length were significantly reduced (Supplemental Figure S2C-E). This suggests possible effects on M-cell function leading to deficits in escape behavior.

Next, we asked if *auts2a* mutants exhibited any deficits in escape behavior. We evoked the C-start escape behavior in partially restrained zebrafish larvae between 6-8 days post fertilization (dpf), by directing a strong jet of water at the otic vesicle (OV) (Figure 2A). The restrained preparation ensures similar location of water jet delivery across larvae. The C-start escape response consists of a large angle contralateral tail bend initiated within 3-13 ms of stimulus delivery (Figure 2B). We measured three parameters associated with the C-start escape response: the probability of initiating escapes (% trials where escapes were observed), the latency (time from stimulus arrival at the OV to movement onset) and the maximum tail bend angle, in wild type, heterozygotes and *auts2a* mutant larvae. First, *auts2a* mutants showed a higher percentage of failures to initiate escape responses compared to heterozygotes and wild type larvae (Figure 2C). While wild type and heterozygote larvae showed a contralateral tail bend response in nearly 98% of trials, *auts2a* mutants were able to generate escapes in only 76% of trials. Mutants also had increased probability of failures (no tail bend response) compared to heterozygote and wild type larvae. Next, we asked if the increased failure rates observed in mutants was due to few individuals that did not respond across any of the trials. Surprisingly, we observed that individual larvae displayed highly variable responses across trials (supplementary video 1). Figure 2D shows responses from 5 wild type and 5 mutant larvae across 6 trials. While all wild type larvae were able to initiate escape responses within 10-20 ms, 4 of 5 *auts2a* mutants showed failures in at least one trial. In addition, the latency to initiate escapes were longer and more variable in mutants compared to wild type (Range: 3-552 ms; Figure 2D, E). Escape response can be generated via multiple pathways. While fast escape responses to head directed stimuli result from activation of the M-cells and its segmental homologs (M-series) (Kohashi and Oda, 2008; Liu and Fetcho, 1999), neural circuitry underlying long-latency escape responses are less well understood (but see (Marquart et al., 2019). Therefore, we next compared latencies of the fast escape response (cut-off latency: ≤20 ms) between wild type, heterozygotes and *auts2a* mutants. Fast escape responses in *auts2a* mutants occur at longer latencies compared to wild type (Figure 2F). Further, the coefficient of variation (CV) of latencies for individual larvae across successive trials was significantly higher for the mutant group in comparison to wild type larvae (Figure 2G). Nevertheless, the kinematics of the escape response were not affected in mutants, as evidenced by the similar maximal tail bend angles observed (Figure 2H). Thus, initiation of tightly regulated, high performance escape response is unreliable and slow in *auts2a* mutants.

**Figure 2.**
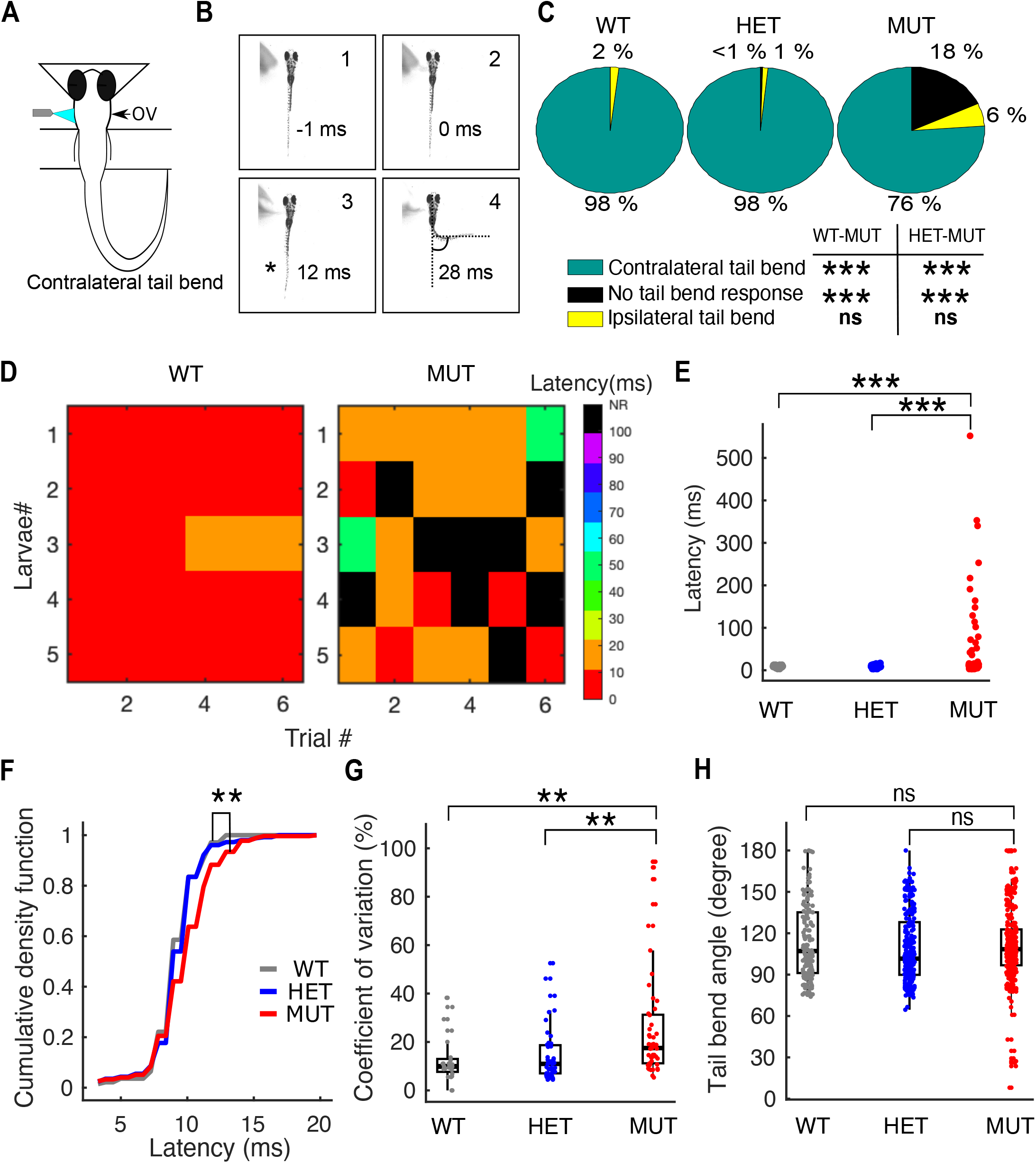
Onset of escape response is delayed and highly variable in *auts2a* mutants. **(A)** Schematic of experimental set up and escape circuit. Escape response was evoked in zebrafish larvae (6-8 dpf) by directing a strong water-jet at the otic vesicle (OV) activating ipsilateral Mauthner (M-cell) and its segmental homologs, MiD2cm and MiD3cm. Escape response is characterised by a large angle tail deflection, contralateral to the direction of water-jet. **(B)** Time lapse of escape response. 1. Pre-stimulus frame. 2. Water jet makes first contact with OV. 3. First visible tail contraction (marked with asterisk). 4. Representative frame showing references used for maximum tail bend angle calculation. **(C)** Pie chart showing percentage of contralateral, ipsilateral and no tail bend responses observed across wild type (n=143 trials), heterozygotes (n= 258 trials) and *auts2a* mutants (n=371 trials). **(D)** Escape latencies across successive trials from five wild type and mutant larvae. Color bar represents escape latencies. NR: no response. **(E)** Comparison of escape response latencies in wild type (WT), heterozygotes (HET) and *auts2a* mutants (MUT). n_WT_=140 trials from 24 larvae, n_HET_=254 trials from 43 larvae and n_MUT_=292 trials from 57 larvae. **(F)** Cumulative density function plot for short-latency escapes (latencies ≤ 20 ms) in wild type (n=140 trials), heterozygotes (254 trials), and *auts2a* mutants (n=273 trials). **(G)** Coefficient of variation of latencies across successive trials in individual larva for wild type (n=24), heterozygotes (n=43) and mutants (n=53) groups. **(H)** Comparison of maximum tail bend angle of contralateral turn between the three groups (n= 140 trials, WT; 254 trials, HET; 281 trials, mutants). Kruskal Wallis; Mann-Whitney test for between-groups comparisons with Bonferroni correction for multiple comparisons. *p<0.025, **p<0.005, ***p<0.0005; ns, not significant.

### Escape response defects in *auts2a* mutants persist on changing the location of sensory stimulation

Stimulation of the OV with water jet activates the M-cell and its homologs, while the same stimulus applied to the tail activates the M-cell alone (Liu and Fetcho, 1999). To investigate if M-cell firing is compromised in *auts2a*^*ncb104*^ larvae, we applied the water jet to the tail at the level of the cloaca (Figure 3A, B). In wild type larvae, tail stimulation with water pulse resulted in fast escape responses with characteristic short latency and contralateral tail bend in 83% trials (Figure 3B, C). Similar to OV stimulation, *auts2a* mutants displayed a high percentage of failures in escape response but similar percentage of ipsilateral tail bend responses to wild type (Figure 3C). Tail stimulation also resulted in significantly longer latencies in mutants than in wild type larvae (Figure 3D). However, no significant difference was observed in the coefficient of variation of latencies across trials in an individual larva between wild type and mutants (Figure 3E), implying that the increased variability of latency seen with OV stimulation could result from recruitment of the M-cell homologs. No difference was observed in tail bend angles between wild type and mutant groups (Figure 3F).

**Figure 3.**
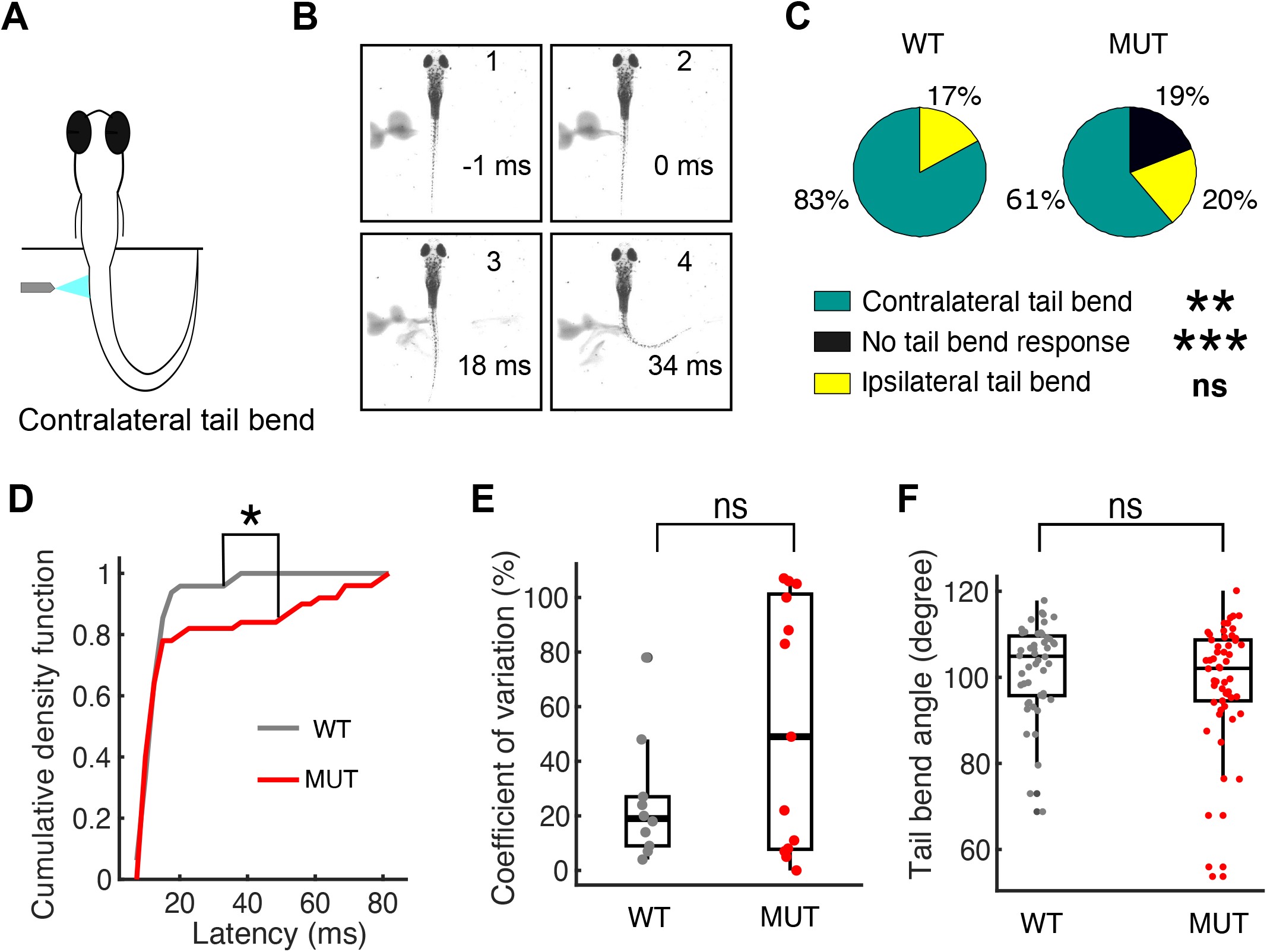
Escape response defects in *auts2a* mutants persist on changing the location of sensory stimulation. **(A)** Schematic of experimental set up and escape circuit. Tail stimulation activates ipsilateral Mauthner but not its homologs and contralateral M-series. **(B)** Time lapse of escape response evoked by tail stimulation. 1. Pre-stimulus frame. 2. Water jet makes first contact with the tail. 3. First visible tail contraction (marked with asterisk). 4. Representative frame for maximum tail bend angle calculation. **(C)** Pie chart showing percentage of contralateral tail bends, ipsilateral tail bends and failures to initiate an escape response between wild type (n=58 trials) and mutants (n=84 trials). **(D)** Comparison of escape response latencies on tail stimulation between wild type (n=48 trials,10 larvae) and *auts2a* mutants (n=50 trials,15 larvae). **(E)** Comparison of coefficient of variation of latencies across successive trials for each larva between wild type (n=10) and mutant (n=13) groups. **(F)** Maximum tail bend angle of turns for WT (n=48 trials) and mutants (n=54 trials). *p<0.05, ***p<0.0001; ns, not significant.

### Mauthner neuron fails to fire reliably in *auts2a* mutants

To directly assess M-cell firing in *auts2a* mutants, we next monitored calcium activity in OGB-1 dextran labelled M-cells upon electrical stimulation of OV (Figure 4A). OV stimulation resulted in a large increase in fluorescence from rest (Figure 4B, C, supplementary video 2) and this response was evoked reliably in wild type larvae (Figure 4B-D). Mutants displayed greater proportion of failures in calcium response (Figure 4B-D, supplementary video 3). Mutants also showed lower peak calcium signals compared to wild type (Figure 4C, E). In addition, the CV of the peak calcium response was not statistically different between wild type and mutant larvae. This implies that the increased variability in latency seen with OV stimulation (Figure 2F) does not arise from variability in response of the M-cell itself.

**Fig 4.**
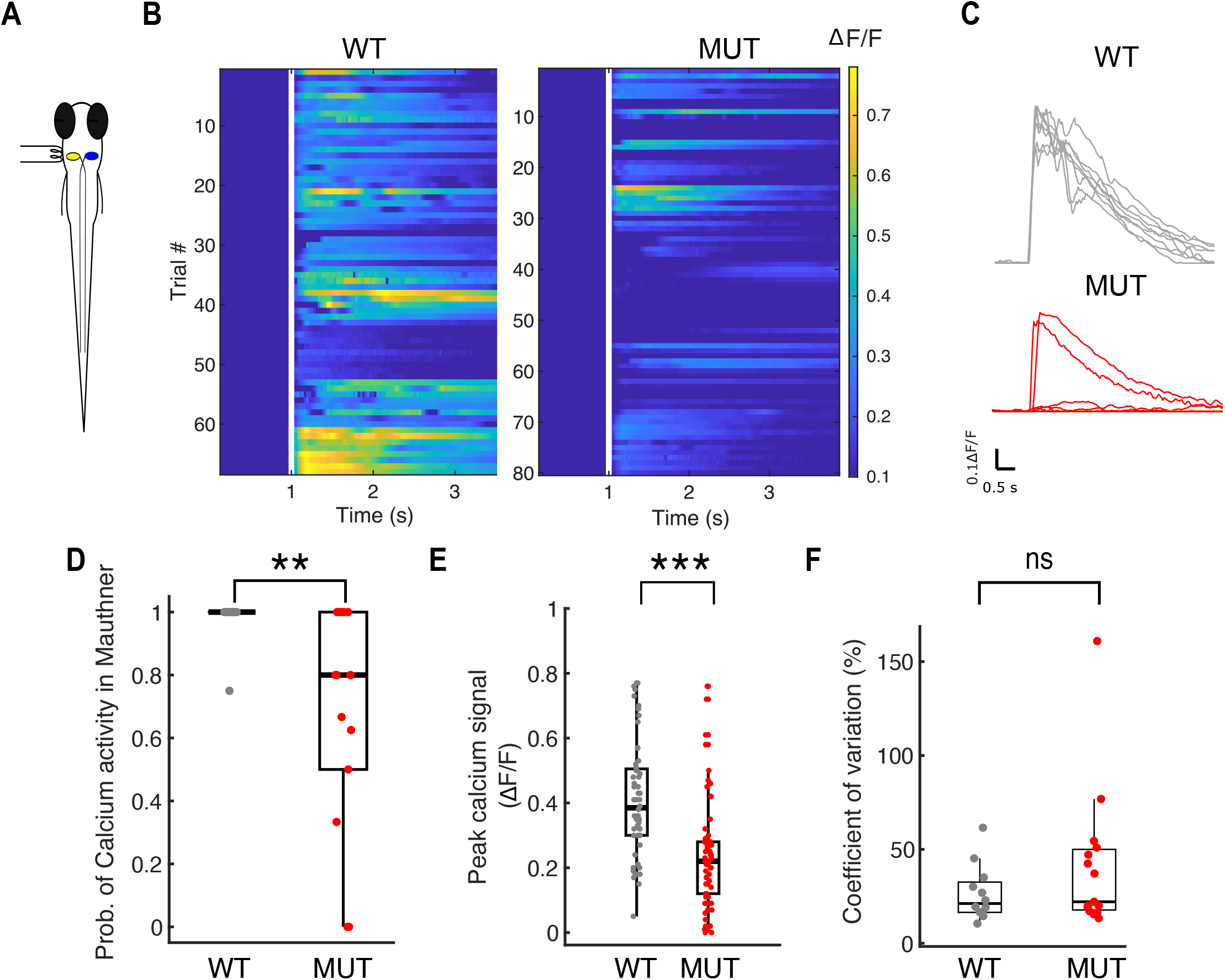
Mauthner neuron fails to fire reliably in *auts2a* mutants. **(A)** Schematic representation of experimental set up. Mauthner neuron was retrogradely labelled with OGB-1 dextran and calcium activity was monitored upon electrical stimulation (40 μA, 1ms) of OV. **(B)** Left : Raster plot of all trials in WT (n=68 trials; 10 larvae) showing consistent calcium activity across several trials on OV stimulation. Right : Calcium responses observed across all trials in the mutant group (80 trials; 14 larvae). White line represents time of stimulus delivery. **(C)** Top: ΔF/F profile of a Mauthner neuron in an example wild type larva across 8 trials in response to electrical stimulation of the OV. Bottom: ΔF/F profile of a Mauthner neuron in an example *auts2a* mutant larva showing sub-threshold response as well large calcium transients across 8 trials upon electrical stimulation of OV. **(D)** Probability of calcium activity response across trials per larva (n_WT_=10 larvae, n_MUT_=14 larvae). **(E)** Peak ΔF/F in WT and mutants (n_WT_=68 trials, n_MUT_= 80 trials). **(F)** Comparison of coefficient of variation of peak ΔF/F between wild type and mutant larvae. Mann-Whitney U test; ***p<0.0001.

### Mauthner neurons in *auts2a* mutants have reduced excitability

Failures in calcium activity response in the mutants after OV stimulation could result either from reduced excitability of M-cells and/or defects in sensory processing involving mechanosensory hair cells and the VIIIth cranial nerve. To ascertain the role of the M-cells, we stimulated its axon (antidromic stimulation) which resulted in calcium activity transients similar to OV stimulation (Figure 5A).

**Fig. 5.**
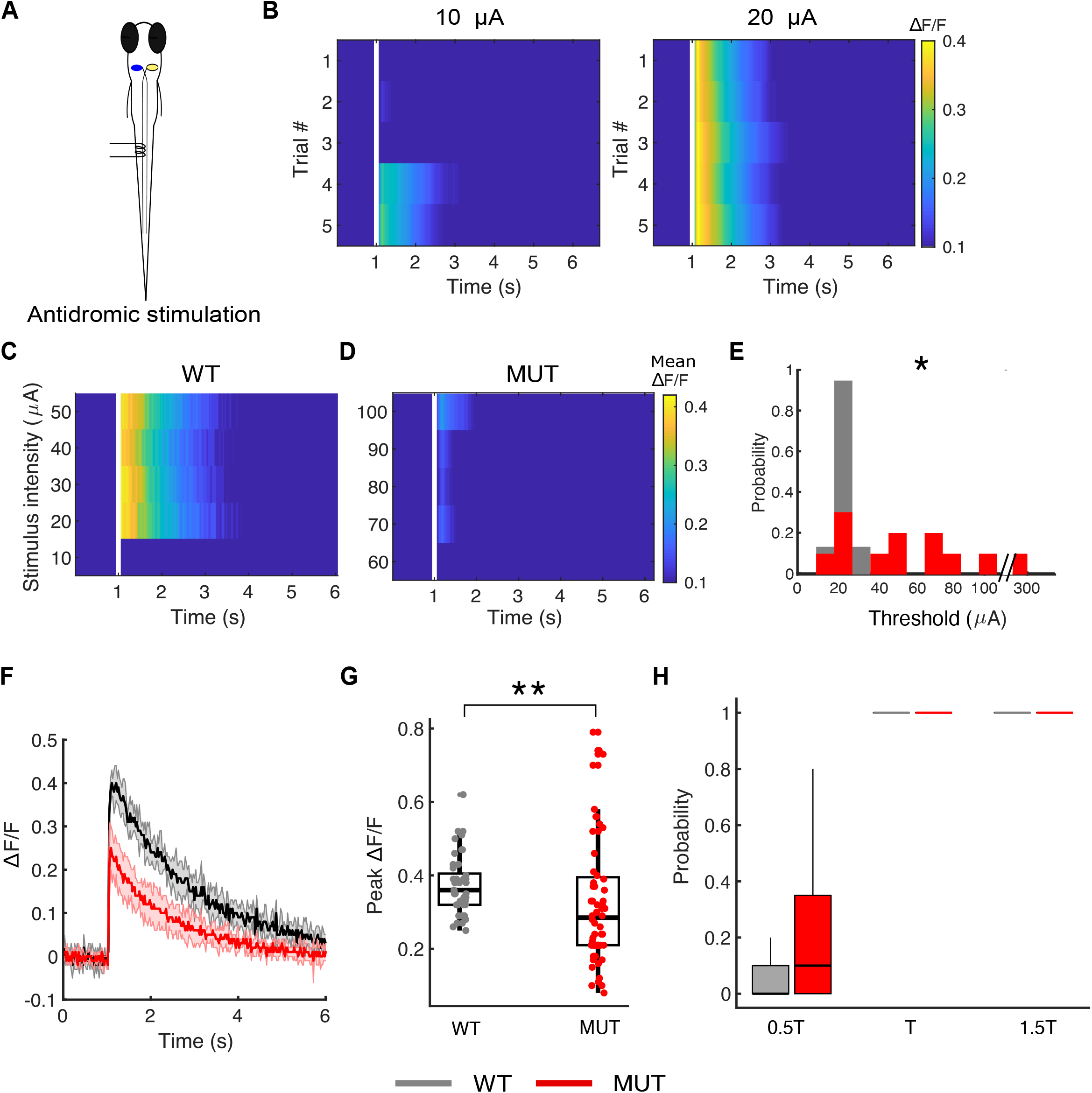
Mauthner neuron in *auts2a* mutants have reduced excitability. **(A)** Schematic of experimental set-up. Calcium activity in Mauthner neuron was observed on antidromic stimulation. Mauthner neuron was retrogradely labelled with OGB-1 dextran. **(B)** ΔF/F profile for an example wild type larva upon antidromic stimulation with 10μA (left) and 20 μA (right) stimulus intensity. Mauthner neuron fired reliably at the threshold intensity of 20 μA. **(C)** Representative raster plot from a wild type larva (left) and mutant larva (**D**) Each row represents average ΔF/F over five trials at respective stimulus intensity. The threshold for calcium activity for wild type larva is 20 μA whereas for the mutant larva is 70 μA. **(E)** Normalised histogram of calcium activity threshold for *auts2a* mutants (n=12 larvae) and WT (n=9 larvae). **(F)** Summary data of probability of calcium activity at 0.5, 1, 1.5 X threshold stimulus intensity for wild type and mutant group. **(G)** ΔF/F profiles for a representative wild type larva (black) and a mutant larva (red) on antidromic stimulation. Shaded regions represents SEM from five trials. **(H)** Summary data of peak calcium signal in wild type (n=45 trials, 9 larvae) and *auts2a* mutants (n=60 trials, 12 larvae). *p<0.05, **p<0.001; Mann-Whitney U test

Due to the large diameter of the M-cell axon, it has the lowest threshold for extracellular stimulation (Kimmel et al., 1982). At low stimulation intensity other cells are unlikely to be activated, making the stimulation M-cell-specific. We defined the threshold intensity as the minimum stimulus intensity at which calcium signals were evoked in the M-cell during all 5 trials (Figure 5B). Compared to wild type, threshold intensity was significantly higher in *auts2a* mutants (Figure 5C-E). However, at intensities equal to or higher than threshold, M-cells responded reliably in both mutants and wild type larvae (Figure 5F) and the peak calcium signal was significantly reduced in mutants compared to wild type larvae (Figure 5G, H), similar to that seen after OV stimulation (Figure 4E, F). These results show that the increased failures, latency and variability in latencies all derive from increased threshold to fire in M-cells. This in turn is due to decreased excitability of these cells in *auts2a* mutants.

## Discussion

We show that Auts2a function is essential for setting the excitability of M-cells in the escape circuit. As a consequence of *auts2a* mutation, reliable firing of M-cells is lost and therefore, escape responses become unreliable also. In trials where neither the M-cell nor its homologs fire, fish fail to generate escape responses, while in trials where only the homologs fire, the latency to respond is longer. *Auts2a* mutant fish exhibit increases in both failures and latencies, indicating deficits in excitability of the M-cell and its homologs (Figure 6). We confirmed the hypoexcitability of the mutant M-cells by directly stimulating the M-axon. Mounting an appropriate and fast escape response reliably is essential for survival and thus Auts2a serves an essential function for the animal. These findings place Auts2a as an important transcriptional regulator of neuronal intrinsic properties, ultimately setting the function of neuronal circuits. This study thus uncovers a hitherto unknown, but essential function of Auts2 family proteins in neural circuit function and behavior.

**Figure 6.**
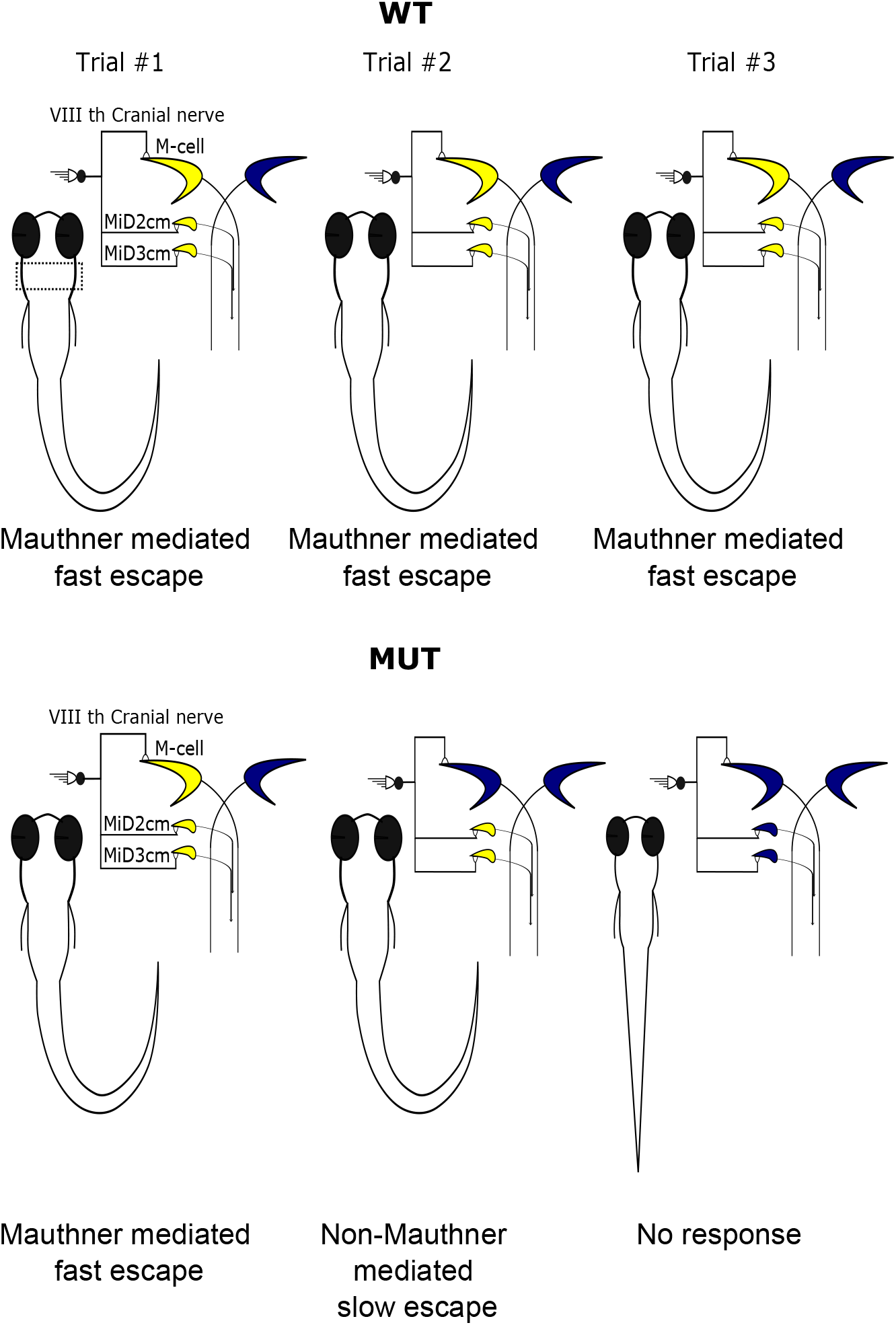
Summary of behavioural abnormalities in escape response in *auts2a* mutants. Top: In response to threatening stimuli ipsilateral Mauthner and its homologs in the hindbrain (marked in dashed box) fire reliably (yellow) resulting in short latency escape responses across consecutive trials in wild type larvae. Bottom: Reduced excitability of Mauthner neurons and homologs in mutants results in decreased probability of firing an action potential and variable behavioral responses (short latency escape response, longer latency non-Mauthner mediated escape response and no response) across consecutive trials.

### The escape circuit as a model circuit for testing gene function

The escape system in zebrafish has been a fantastic tool for dissecting the genetic underpinnings of behavior, learning and decision-making (Gahtan and Baier, 2004; Wolman and Granato, 2012). Forward genetic screens identified mutants with specific deficits in generating escapes such as the *twitch twice* and *space cadet* mutants (Burgess et al., 2009; Granato et al., 1996; Lorent et al., 2001). More recent studies have reported mutants with deficits in sensitivity (Marsden et al., 2018), habituation (Wolman et al., 2015), pre-pulse inhibition (Burgess and Granato, 2007) or in deciding between Mauthner-mediated short latency escapes and non-Mauthner mediated long latency escapes (Jain et al., 2018). The specific defects identified in these studies range from errors in axon guidance, extracellular calcium sensing and IGF signaling in M-cells and other members of the escape circuit. Our study makes an important addition to these studies by identifying Auts2a to be a direct genetic determinant of excitability in M-cells and its homologs.

M-cells are specified soon after gastrulation and evoke escape responses to touch in larvae as young as early as 2 dpf (Kimmel et al., 1974; Kohashi et al., 2012). As the larvae mature, M-cells drive startle behaviors in response to auditory/vestibular stimulation as well. Concomitant with these changes, the firing behavior of M-cells changes from firing multiple action potentials upon reaching threshold to firing only a single action potential after 4 dpf. This change in firing behavior of the M-cells drives maturation of the escape behavior from one involving multiple C-bends to that with only a single C-bend followed by routine swimming. The alteration in M-cell firing behavior is in part due to the expression of distinct types of potassium channels including those that are sensitive to dendrotoxin (Watanabe et al., 2013, 2017). These studies underline the critical importance of regulating the intrinsic properties of M-cells for generating appropriate C-start behaviors. Hyperexcitability will result in multiple C-starts while hypo-excitability in M-cells and its homologs, as seen in *auts2a* mutants will lead to failures in escapes.

### Mechanism of action

ChIP-seq analysis revealed that in mouse brain, AUTS2 binds preferentially to promoter and enhancer regions of genes involved in nervous system development (Gao et al., 2014; Oksenberg et al., 2014). In mouse forebrain alone, 784 AUTS2 binding sites were in promoter regions and 1146 sites were distal to promoter regions. Regardless of whether the sites were within promoter regions, these AUTS2 binding sites were found to be associated with genes implicated in ASDs. Further, binding of AUTS2 to these sites seems to activate their expression resulting in higher transcript levels. Importantly, among the targets of AUTS2 binding are genes associated with intracellular calcium homeostasis such as pumps and transporters, voltage-gated calcium channels and sodium channels, potassium channels as well as synaptic receptors (Oksenberg et al., 2014). One of the targets of AUTS2 binding, the voltage-gated calcium channel subunit cacna2d2 is known to regulate the calcium current density and the activation/inactivation kinetics of P/Q type, L-type, N-type and T-type calcium channels. Mutations in voltage-dependent calcium channels are known to associate with ASDs (Krey and Dolmetsch, 2007; Heyes et al., 2015; Lory et al., 2020). Loss of Auts2a in zebrafish could potentially reduce M-cell excitability via dysregulation of intracellular calcium homeostasis and by directly affecting calcium channel function during depolarization. Another important pair of targets of AUTS2 binding are Kcnip1 and Kcnip4, which are potassium channel interacting proteins (Oksenberg et al., 2014). These proteins contain an EF-hand domain, sense intracellular calcium and regulate A-type transient potassium currents. A-type potassium channels are critical determinants of firing threshold and latency. Lastly, Auts2 binding is also seen within genes coding for a number of voltage-gated potassium channels (Oksenberg et al., 2014). Thus, loss of Auts2a might interfere with expression levels of one or more of these targets leading to hypoexcitability of M-cells. Wherever it is expressed, Auts2 might be essential for determining neuronal excitability by setting levels of expression of ion channel genes or their regulatory partners. Manipulations that reduce global activity levels of neurons in culture lead to homeostatic resetting of intrinsic and synaptic properties (Turrigiano et al., 1998; Desai, 2003; Turrigiano and Nelson, 2004), a process that requires transcription (Ibata et al., 2008). Recently, it was shown that manipulations that induce homeostatic plasticity also trigger significant upregulation of AUTS2 expression (Schaukowitch et al., 2017). Thus, on the basis of our study and these earlier studies, we posit that Auts2 is important for setting and maintaining the excitability set-points of neurons.

### Alterations in neuronal excitability and ASDs

Since ASDs are frequently also comorbid with epilepsy, and because mutations that reduce inhibition or increase excitation frequently associate with them, ASDs were initially thought to result from hyperexcitation in the central nervous system (Rubenstein and Merzenich, 2003). However, it is becoming clear that balance between excitation and inhibition is key and when that balance is tilted towards more excitation or more inhibition, network function is impaired leading to ASD-like phenotypes (Nelson and Valakh, 2015; Lee et al., 2017; Sohal and Rubenstein, 2019). Activity imaging using expression of the immediate early gene *c-Fos* reveals hypoactivity in much of the forebrain in a mouse model of Rett syndrome (Kron et al., 2012). Reduced connectivity and reduced activation of forebrain structures at rest have also been reported in fMRI studies of human autistic subjects (Minshew and Keller, 2010). These studies underline the fact that since ASDs are associated with mutations in multiple genetic pathways, they should be thought of as diseases resulting from abnormal excitation to inhibition balance. Consistent with this view, a recent study demonstrates that in *Auts2* mutant mice, hippocampal pyramidal neurons receive increased excitatory synaptic inputs with no change in the amount of inhibition received, upsetting the excitation to inhibition balance (Hori et al., 2020). We have shown that loss of function of Auts2a in zebrafish leads to hypoexcitability in an identified neuron, the M-cell, which is responsible for driving fast escapes. Our identification of this specific deficit in an identified neuron in a circuit with defined behavioral function is significant as it pinpoints to specific cellular mechanisms due to which excitation to inhibition balance is altered in the autistic brain. Taking these studies together, *Auts2/Auts2a* might play circuit specific functions in maintaining the excitation to inhibition balance by triggering transcriptional programs for the expression of specific combinations of ion channels and synaptic receptors.

## Acknowledgements

The authors would like to thank the following sources of funding support: Wellcome Trust-DBT India Alliance Intermediate and Senior fellowships (VT), Department of Biotechnology (VT), Science and Engineering Research Board, Department of Science and Technology (VT), Department of Atomic Energy (VT-12-R&D-TFR-5.04-0800), CSIR-UGC fellowship (UJ), NCBS Career Development Fellowship (IK). VK’s laboratory in the IMCB, Singapore was supported by the institutional grant to the IMCB by the Agency for Science, Technology and Research of Singapore. VK in Poland was supported by the Opus grant of the National Science Foundation (NCN), Poland (2016/21/B/NZ3/00354).

The authors would also like to thank T.P. Jagadeesh for maintaining the fish facility.

## Author Contributions

U.J., I.K. and V.T., conception and design; U.J. acquisition and analysis of behavior and imaging data; I.K., generation of the mutant lines; V.K., and V.T., funding acquisition and resources; U.J., I.K. and V.T., drafting the manuscript.

## Materials and Methods

### Fish care and use

Zebrafish (*Danio rerio*) of AB strain and Indian wild type were housed in aquarium tanks at 28.5°C with a 14:10 hours light:dark cycle. Fish were maintained according to established protocols as previously described (Westerfield, 2000) in agreement with the Institutional Animal Ethics Committee and the Institutional Biosafety Committee, National Centre for Biological Sciences. For fin amputation, fish were briefly anaesthetized in 0.01% Tricaine (MS-222, Sigma-Aldrich), the caudal fin was cut and fish were immediately returned to fresh water. Experiments were performed on 6-8 days post fertilization (dpf) larval zebrafish at room temperature. Larvae were maintained in 14:10 light-dark cycle at 28°C in E3 medium (5 mM NaCl, 0.17 mM KCl, 0.33 mM CaCl_2_ and 0.33 mM MgSO_4_, pH 7.8). Larvae were treated with 0.003% 1-phenyl-2-thiourea in 10% Hank’s saline at 24 hpf to remove pigments, for calcium imaging experiments.

### TALEN design, construction and synthesis

TALENs specific to *auts2a* were manually designed using criteria as described previously (Bedell et al., 2012) and assembled according to the established protocol (Sanjana et al., 2012). The plasmid kit used for building TALENs was a gift from Dr. Feng Zhang (Addgene kit #1000000019). Once assembled into a destination vector, TALENs then were re-cloned into pTNT vector (Promega) containing synthetic polyA tail and T7 terminator sequence. The *auts2a* TALEN recognition sequences are as follows: left TALEN, CCAGCTGGGAGTGCCT and right TALEN, GTAATAGCACTTTAGGTGG. Between the two binding sites is a 16-nt spacer with a *Kpn*I site (ACTCA<u>Ggtacc</u>agtca, *Kpn*I site is underlined, intronic sequence is in lower case), facilitating identification of mutations by PCR and KpnI restriction digestion. The spacer region overlaps with the donor splice site at exon 8 (Figure 1A). Capped mRNA was synthesized by *in vitro* transcription using mMESSAGE mMACHINE T7 kit (Invitrogen) and purified with RNeasy Mini kit (Qiagen). Zebrafish embryos at the 1-cell stage were microinjected with 200 pg RNA (100 pg each of left and right TALEN mRNA). At such a dose, over 70% embryos survived and showed TALEN-induced somatic *auts2a* gene modifications.

### DNA isolation and genotyping

Genomic DNA was isolated from either embryos or fin clips using HotSHOT method (Meeker et al., 2007). One microliter solution was then used in a 25⎧L PCR containing the following reagents at these concentrations: 200 nM each gene-specific primers (forward 5’-TCAGCGAACCCTACAGCTTCACACA-3’ and reverse 5’-TGGGGTACGCACCATGGGCGGTGCA-3’), 0.2 mM dNTPs, 1xPCR buffer and 0.625 units One*Taq* HotStart DNA polymerase (New England BioLabs). Reaction was amplified using the following conditions: 94°C for 1 min; 40 cycles of 94°C for 20 s, 68°C for 1 min; followed by 68°C for 1 min. PCR products were purified using PCR purification kit (Qiagen) and digested with restriction enzyme *Kpn*I-HF (New England BioLabs) at 37°C for 60 min. The resulting reactions were loaded onto a 1.8% agarose gel and electrophoresed in 1x Tris-acetate-EDTA (TAE) buffer. Mutations were assessed by loss of restriction enzyme digestion. To verify mutations, the gel purified uncut PCR products were cloned into pCRII-TOPO vector (TOPO TA Cloning kit, Invitrogen) and sequenced.

### RT-PCR and RNA isolation

Total RNA was isolated from wild type and *auts2a*^*ncb104*^ heterozygote and homozygote embryos using an RNeasy Mini kit (QIAGEN), and first-strand cDNA was synthesized from 1 ⎧g of total RNA by oligo(dT) priming using SMARTScribe Reverse Transcriptase (Clontech) according to the manufacturer’s protocol. Amplification of cDNA was performed using Herculase II Fusion DNA polymerase (Agilent). Identity of amplified PCR products was verified by direct sequencing.

### Whole-mount Immunohistochemistry

Embryos were fixed in 4% PFA at 4°C overnight, washed 3 times for 15 min in PBST (1 x PBS, 0.1% Tween-20) and permeabilized in 0.1% Triton X-100 in 0.1% sodium citrate for 30 min at 4°C. Then embryos were incubated for 2 hours in 5% Blocking Reagent (Roche) in MAB (150 mM maleic acid, pH 7.5, 100 mM NaCl, 0.1% Tween-20) at room temperature. Embryos were incubated with 3A10 antibodies (DSHB, 1:200) overnight at 4°C, washed 4 times for 30 min in MAB and incubated with HRP-conjugated goat anti-mouse F(ab)_2_ fragments (Molecular Probe, 1:500) for 6 hours at room temperature or overnight at 4°C. Embryos were extensively washed in PBST, stained with 3,3’-diaminobenzidine (DAB) and washed several times in PBST. Embryos were kept in 50% glycerol in PBS at 4°C until further imaging.

### Head restrained preparation

For behaviour and calcium imaging experiments, larvae were embedded in 2% low gelling agarose (Sigma-Aldrich, Missouri, USA). E3 medium was added after the agarose congealed. Agarose around the tail and the otic vesicle (OV) were removed for observing tail movements and application of water pulse for behavioural experiments.

### High-speed recording of escape response

Escape responses were evoked by applying a water pulse to the OV or to the tail at the level of cloaca, with a glass capillary (tip diameter 0.05-0.06 mm) mounted on a micromanipulator (Narishige; Tokyo, Japan). The water pulses were generated by a pressure pulse of 10 ms duration and pressure of 30 psi from microinjection dispense system (Picospritzer III, Parker Hannifin, Ohio, USA). Videos were acquired at 1000 fps with a high speed camera (Phantom Miro eX4, Vision research, New Jersey, USA) mounted on a stereo microscope (SZX16, Olympus, Tokyo, Japan) at 512 × 512 pixel resolution and 500 μs exposure time. Methylene blue (1%) was used to visualize the water jet for tracking its contact with the larva. Six trials (three on each side) were performed on each larva.

### Retrograde labelling of Mauthner neurons

Mauthner neurons were retrogradely labelled with fluorescent calcium indicator Oregon Green Bapta-1 dextran, 10000 mW (Invitrogen, California, USA). Larvae were first anaesthetised in 0.01% MS-222 (Sigma Aldrich; Missouri,USA) in E3 medium. 25% OGB-1 in 10% Hank’s Balanced Salt Solution (HBSS; 137 mM NaCl, 5.4 mM KCl, 0.25 mM Na_2_HPO_4_, 0.44 mM KH_2_PO_4_, 1.3 mM CaCl_2_, 1.0 mM MgSO_4_, 4.2 mM NaHCO_3_) was pressure injected with a glass microcapillary into the spinal cord (at the level of cloaca) using a Picospritzer. Post injection, larvae were allowed to recover in HBSS for >12h.

### Electrical stimulation

Electric shock stimuli (40 μA, 1 ms) were delivered using a bipolar electrode (FHC, Bowdoin, ME, USA) placed at the OV. The pulse was generated using ISO-Flex stimulus isolator (A.M.P.I., Jerusalem, Israel), triggered by pClamp (Molecular devices, California, USA). For antidromic stimulation of the M-axon, larvae were anaesthetized in 0.01% MS-222 (Sigma-Aldrich; Missouri, USA) and were pinned down through notochord using fine tungsten wire (California Fine Wire). The MS222 was then replaced by external solution (composition: 134 mM NaCl, 2.9 mM KCl, 1.2 mM MgCl_2_, 10 mM HEPES, 10 mM glucose, 2.1mM CaCl_2_, 0.01 mM D-tubocurarin; pH 7.8; 290 mOsm) and skin along the tail was carefully removed using forceps (Fine Science Tools, Foster City, USA). Muscles in a hemi-segment (between the 10^th^ and 13^th^ myotomes) were carefully removed to expose the spinal cord. The bipolar electrode was placed on top of the exposed spinal segment and brief electrical stimuli of increasing strengths (10 μA onwards, 1 ms in duration) were delivered.

### Calcium imaging

Calcium activity upon electrical stimulation of OV/M-axon, in retrogradely labelled Mauthner neurons, was imaged at 35-45 frames per second using an EMCCD camera (Evolve, Photometrics, UK) mounted on a compound microscope (BX61W1, Olympus, Tokyo, Japan) with a water-immersion objective (LUMPlanFL 60X) and Image-Pro Plus (Media Cybernetics, UK) acquisition software.

### Analysis

Data were analysed using MATLAB (Mathworks; Massachusetts, USA) and Fiji (NIH).

### Behavioural analysis

Latency was defined as the time taken from when the water jet made contact with the larva to the first visible tail contraction. Tail bend angle was calculated by measuring the angle formed by joining a straight line passing through the tail at maximum bend and a straight line passing through the head and the tail in a pre-stimulus frame (Figure 2C). Escape responses were defined as contralateral tail bends with latency ≤100 ms (Kimmel et al., 1974; Kohashi and Oda, 2008). Tail bend responses with latency >100 ms or no observable tail bend upto 1 s post stimulus delivery were classified as failures (‘no response’).

### Calcium activity analysis

Relative changes in fluorescence from resting (ΔF/F) were calculated post background and movement correction. Activity in Mauthner neurons was considered to have occurred only for ΔF/F traces with minimum peak ΔF/F of 0.1 and full width at half maximum of at least 1 s. Threshold for antidromic stimulation of Mauthner was defined as the minimum stimulus intensity at which the Mauthner neuron showed calcium activity for all trials at the given stimulus intensity.

**Dendritic length** and **soma volume** of Mauthner neurons were measured using simple neurite tracer and volume viewer plugins in Fiji (Schindelin et al., 2012).

### Statistics

Data were tested for normality using one sample Kolmogorov-Smirnov test (p<0.05) and equality of variance with *F* test (p<0.05). Two sample t-test or Mann-Whitney U test were performed for comparisons between two groups and Kruskal-Wallis test was used for comparing three groups. Chi-square test was performed for comparison of proportions.

