## Supplemental Figure 1 for "High behavioural variability mediated by altered neuronal excitability in *auts2* mutant zebrafish"

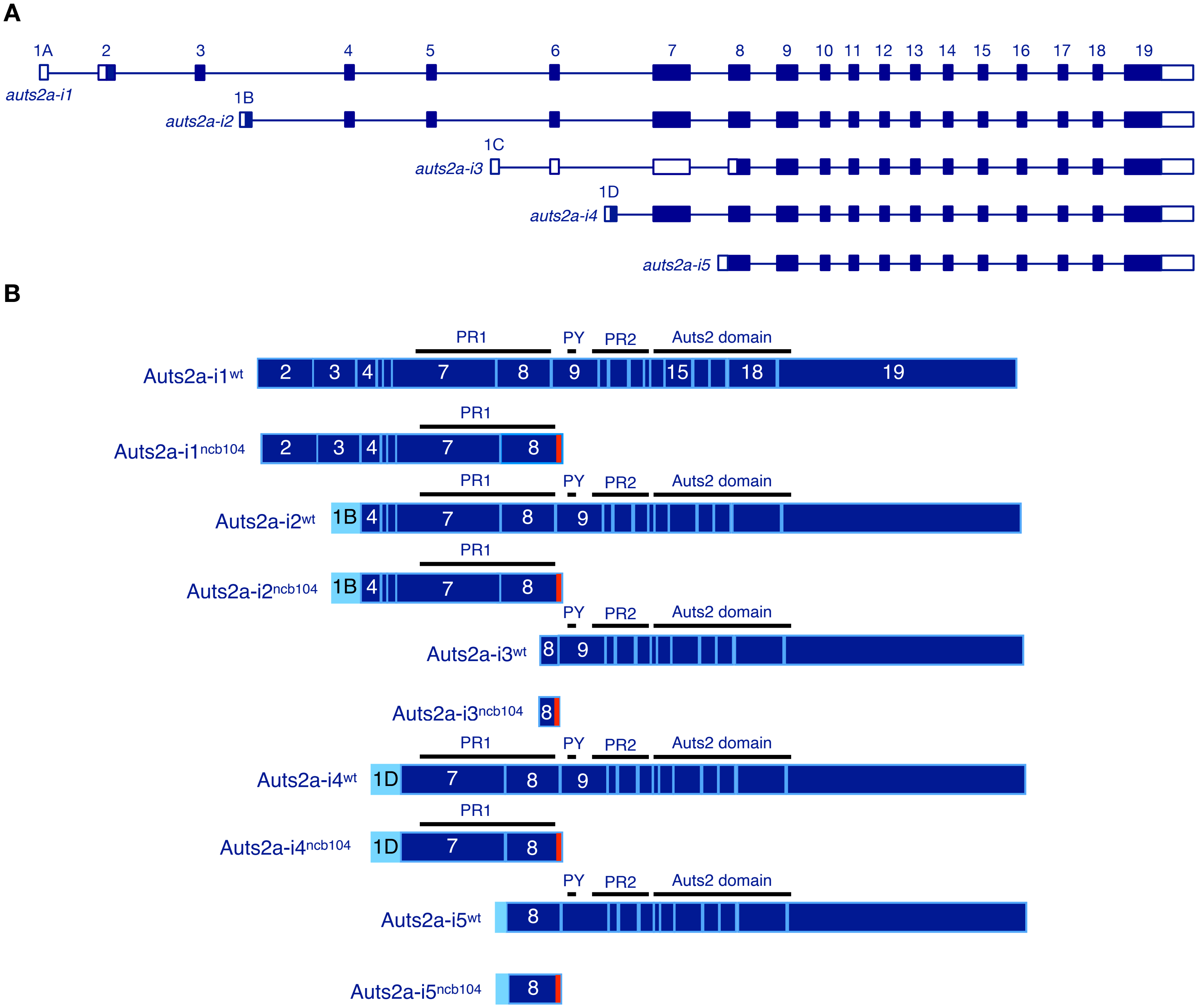


**Figure S1.** ***Auts2a*** $^{\mathit{ncb104}}$ **allele in different** ***auts2a*** **isoforms.**

**A.** Overview of transcripts generated from the *auts2a* gene (modified from Kondrychyn et al. 2017). Noncoding and coding exons are depicted as open and filled bars, respectively. Alternative transcription start sites are used to generate *auts2a* isoforms. **B.** Schematic structure of Auts2a$^{wt}$ and Auts2a$^{ncb104}$ proteins translated from the different *auts2a* isoforms. Positions of coding exons are marked for the reference (relative exon size is not in scale). Missense amino acids preceding premature stop codon are shown in red. Exons 1B and 1D code the alternative N-terminal amino acids. PR1 spans exons 7 and 8, PR2 spans exons 9-13 and the Auts2 family domain spans exons 14-19.
