## Supplemental Figure 2 for "High behavioural variability mediated by altered neuronal excitability in *auts2* mutant zebrafish"

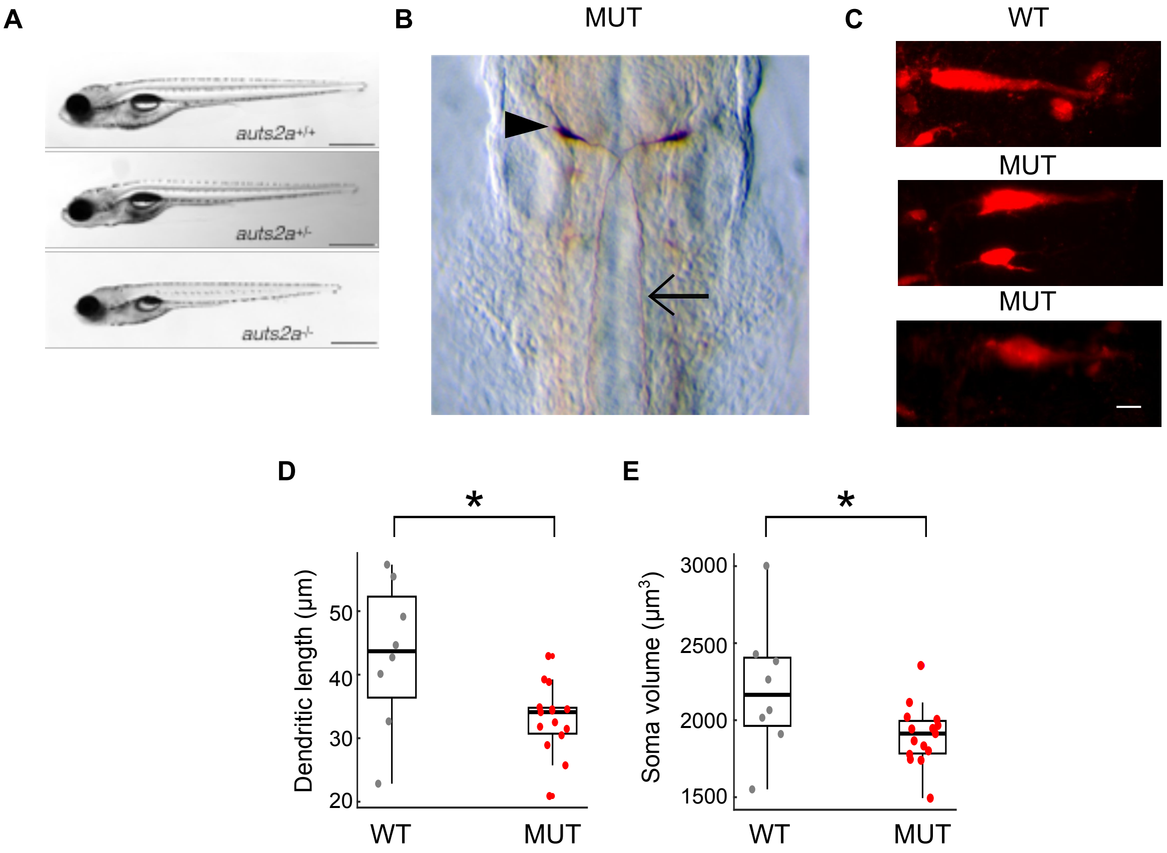


**Figure S2. Morphological characterization of *auts2a* mutants.** **(A)** Bright field images of wild type, auts2a$^{ncb104}$ heterozygote and homozygote larvae. Scale bar represents 2 mm **(B)** Whole mount immunostaining with 3A10 antibody at 30 hours post fertilization (hpf) in *auts2a* mutants. Arrowhead points to the cell body of Mauthner neuron and arrow points to the axon. **(C)** Z-projection of Mauthner neuron in a WT larva (top) and two mutant larvae (middle, bottom). Mauthner neurons were reterogradely labelled with tetra-methyl rhodamine dextran (red) **(D)** Comparison of length of lateral dendrite in Mauthner neuron **(E)** Comparison of soma volume of Mauthner neuron in WT and mutant larvae. Scale bar represents 10µm. $n_{WT}$=8 cells from 7 larvae; $n_{MUT}$=15 cells from 12 larvae. *, $p<0.05$; Mann-Whitney U test.
